# Duet model unifies diverse neuroscience experimental findings on predictive coding

**DOI:** 10.1101/2025.07.12.664417

**Authors:** John H. Meng, Jordan M. Ross, Jordan P. Hamm, Xiao-Jing Wang

## Abstract

The brain continuously generates predictions about the external world. When stimulus X is presented repeatedly, the brain predicts that the next one is also X. A deviant stimulus Y elicits a stronger sensory response than the baseline, reflecting the amplification of an unexpected stimulus. Here, we introduce the duet predictive coding model, a minimal and biologically plausible framework in which neurons encode both positive and negative prediction errors. This model reproduces neural responses observed in vision and audition across diverse predictive coding paradigms, particularly omission. Our proposed circuit mechanism predicts (1) neurons tuned to negative prediction errors in the oddball paradigm, supported by experimental evidence in mice; (2) the magnitude of unexpected responses quantitatively depends on the dissimilarity between standard and deviant stimuli and diminishes with increasing interstimulus interval. Our findings suggest that the brain’s deviance detection relies on dual-error computation, offering a unifying explanation across seemingly disparate experimental protocols.

## Introduction

In our daily lives, we consciously or unconsciously predict what will happen around us. We generate internal expectations and compare them with incoming stimuli, such as a sound or a visual pattern (Figure 1A). For example, when playing the piano, we can often predict the upcoming note and may disengage our attention from it. Consequently, an off-key note may catch us by surprise. Similarly, we can detect an out-of-tune sound during a live classical performance if we are familiar with the melody. In a simplified scenario, when our internal expectations and incoming stimuli align, we tend to ignore the sensory input (Figure 1B). In contrast, when expectations and stimuli do not match, the brain may generate a deviance-related signal, enabling us to process the unexpected event with heightened vigilance or urgency (Figure 1C). This deviance signal may arise in two forms: when a stimulus occurs without a corresponding internal prediction, like an uninvited guest showing up, or when an expected stimulus fails to occur, as in the case of a speaker not showing up to a seminar. Both types of errors have been detected at the sensory areas [16, 17, 28, 27].

**Fig. 1.**
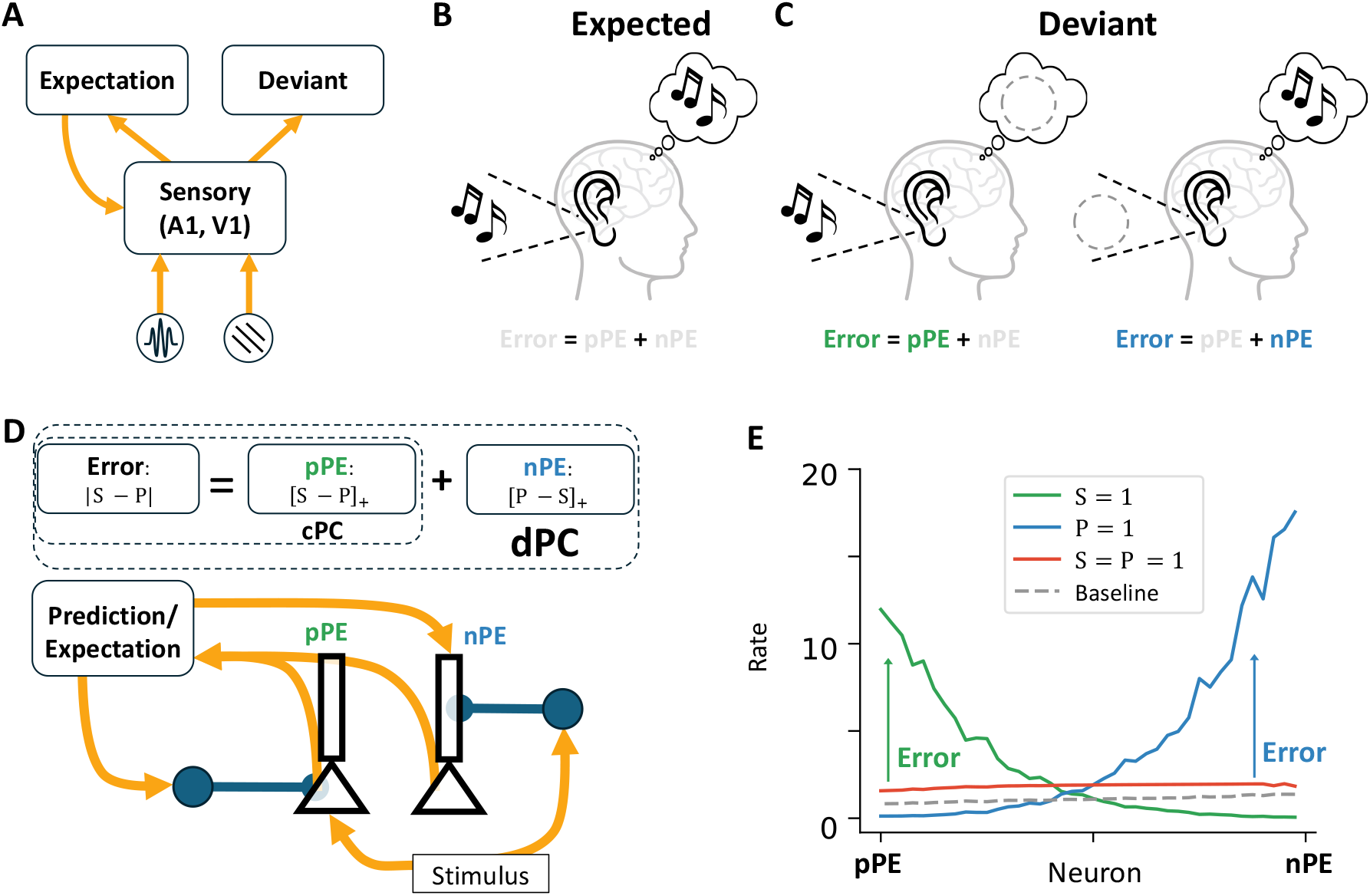
Concepts of duet predictive coding (dPC). (A) Illustration of deviant computation by comparing the expectation with different types of incoming stimuli. (B) An expected case when the stimulus matches the internal expectation. (C) Deviant cases where (Left) stimulus is received without a corresponding expectation, or (Right) an expected stimulus is omitted. (D) Conceptual necessity in error calculation. Classic predictive coding (cPC) only considers positive prediction error (pPE), while dPC accounts for both pPE and negative prediction error (nPE). The pPE neurons are excited by the stimulus (S) and inhibited by the prediction (P) through local inhibitory neurons, encoding the positive error ([S *−* P]_+_). Similarly, the nPE neurons are excited by the prediction and inhibited by the stimulus, thereby encoding the negative error ([P − S]_+_). The required connectivity can be reached through local, biologically feasible learning rules [30, 20]. (E) Firing rates of neurons in different contexts after learning based on [30]. The overall firing rate is close to baseline when stimulus and prediction match. Importantly, in dPC, the prediction input is net excitatory when presented in isolation (blue vs baseline) but becomes net inhibitory when paired with the stimulus (red vs green).

The mathematical formulation of deviance detection can be simplified as the computation of the absolute difference (error) between the incoming stimulus and the internal prediction signal (Figure 1D). This error can be split into two components: the positive prediction error (pPE), representing the positive part of the absolute difference ([S−P]_+_), and the negative prediction error (nPE), representing the remaining negative part ([P − S]_+_).

Since neurons in sensory areas typically have low baseline firing rates [8, 34, 39], it is challenging for a single group to encode bidirectional changes, which are required for representing error. Historically, the classic predictive coding framework (cPC), developed by Rao and Ballard [38], focuses on the pPE component in explaining the reduced visual response to the same stimulus when it is expected. The cPC model has also been used to account for mismatch negativity (MMN) observed in fMRI and electroencephalography (EEG) [15, 52]. Neglecting the nPE component leads to the misconception that the top-down input in the predictive coding must be inhibitory [4]. However, top-down inputs can also be excitatory [58, 10, 23, 57], a fact often cited to challenge the generality of the predictive coding framework [11, 32, 22].

In addition, omitting the nPE component limits cPC’s capacity to account for omission-evoked responses [18, 28, 27]. In such paradigms, subjects build internal predictions through repeated stimulus presentation, followed by an unexpected omission. During the omission, sensory cortices exhibit robust responses, reflecting an error signal between the internal expectation and the absence of the stimulus (Figure 1C, right), which is difficult to explain solely by considering the pPE component.

Instead, we propose that the duet model of predictive coding (dPC), which includes the nPE and pPE components, accounts for the top-down excitation connection. The core idea of dPC is that neurons remain near baseline activity when stimulus and prediction co-occur, as in expected conditions (Figure 1E). When either input is presented alone, excitation in the driven population outweighs inhibition in the suppressed population, allowing the population to act as an effective error detector (Figure 1E). In contrast to cPC, the overall response of prediction input from the local sensory circuit is net excitatory in isolation, which agrees with the observed excitatory response from the top-down signal [4, 22] (Figure E, blue vs baseline). This response becomes net inhibitory when paired with a stimulus (Figure E, red vs. green), consistent with cPC’s central idea that responses to expected stimuli are suppressed. Our previous work showed that pPE and nPE neurons can co-emerge within a circuit via a local learning rule [30], explaining a broad range of motor-sensory phenomena under the dPC framework [1, 24, 3]. Similar connectivity can be learned by enforcing excitation–inhibition balance at both the soma and dendrite of pyramidal cells [20].

Conceptually, including nPE neurons is counterintuitive because they encode the rectified difference between prediction and stimulus ([P − S]_+_), which means some sensory-cortical neurons reduce their activity in response to the stimulus. Nevertheless, recent studies have observed nPE neurons in primary visual cortex (V1) during motion–visual mismatch [1, 24], audio–visual mismatch [13], and in auditory cortex during motor–auditory mismatch [46]. In these tasks, the self-generated motor signal acts as the prediction, compared against sensory input that either results from self-motion (closed-loop) or not (open-loop). Negative prediction errors arise when expected stimuli are interrupted during movement, such as when visual flow is paused while the animal runs on a treadmill in a virtual environment [1]. In such cases, population responses increase, and neurons that respond selectively to negative errors are identified as nPEs. Notably, these neurons also likely exhibit a shared transcriptomic profile that distinguishes them from pPE neurons [35].

However, the prediction signal is often considered an internally generated signal independent of motor output. In such cases, it arises from stimulus history rather than movement. For example, in classic MMN studies [48], EEG responses were stronger to rare tones than to frequent ones, when subjects remained still. Here, the prediction signal is thought to originate from prefrontal or other higher-order cortical areas [26, 15, 22]. If the transcriptomic subtype enriched for nPE neurons in motor-sensory tasks [35] is also present in these contexts, do they still function as nPE neurons when predictions are history-based? If so, how can they be identified and distinguished from pPE neurons?

In this work, we systematically investigate the role of nPE neurons under the dPC framework across diverse predictive coding paradigms using our model and analysis. We first show that the dPC framework explains omission responses via the nPE component [28, 27], and aligns with MMN literature [14]. We then demonstrate how the model captures single-neuron responses in oddball paradigms, predicting a joint contribution of pPE and nPE neurons during deviance detection. Finally, our theory predicts distinct behavioral signatures that differentiate pPE and nPE neurons, which we validate through analyzing experimental data.

## Results

### dPC reproduces omission response

Recently, studies by [28, 27] demonstrated that omission responses can be detected in single neurons in the primary auditory cortex (A1). In their experiment, a sound was presented repetitively at fixed time intervals, with occasional omissions—trials in which a sound was unexpectedly absent. Recordings from the mouse A1 revealed that some neurons consistently responded to these omissions. The authors hypothesized that mice formed an internal expectation of the upcoming sound with precise temporal resolution. When a predicted sound failed to occur, this internally generated prediction signal reached the auditory cortex and triggered the omission response. This suggests the top-down input excites the sensory area, which may not be compatible with the cPC framework.

We investigated this omission protocol within our dPC framework (Figure 2A). We used the learned local connectivity from our previous work [30] as the synaptic architecture. To simulate internal prediction originating from a high hierarchical area, we introduced a leaky integrator acting as a predictor, which generates the top-down prediction input (see Methods for details). This predictor receives bottom-up input uniformly from all neurons in the sensory area and operates on a slower timescale [6, 44] than individual sensory neurons. The top-down prediction signal is temporally aligned with the stimulus window, an essential feature for reproducing omission responses observed at precise expected time points in the experimental data. In our model, this temporal alignment is implemented via a binary timing mechanism for simplicity. The precise timing can be achieved through spike-timing-dependent plasticity, as proposed in [52]. The stimulus is delivered at fixed intervals and adapted across repetitions. After a set number of repetitions (*N*_rep_ = 6 in this case), the stimulus is omitted, simulating an ‘omission’ trial.

**Fig. 2.**
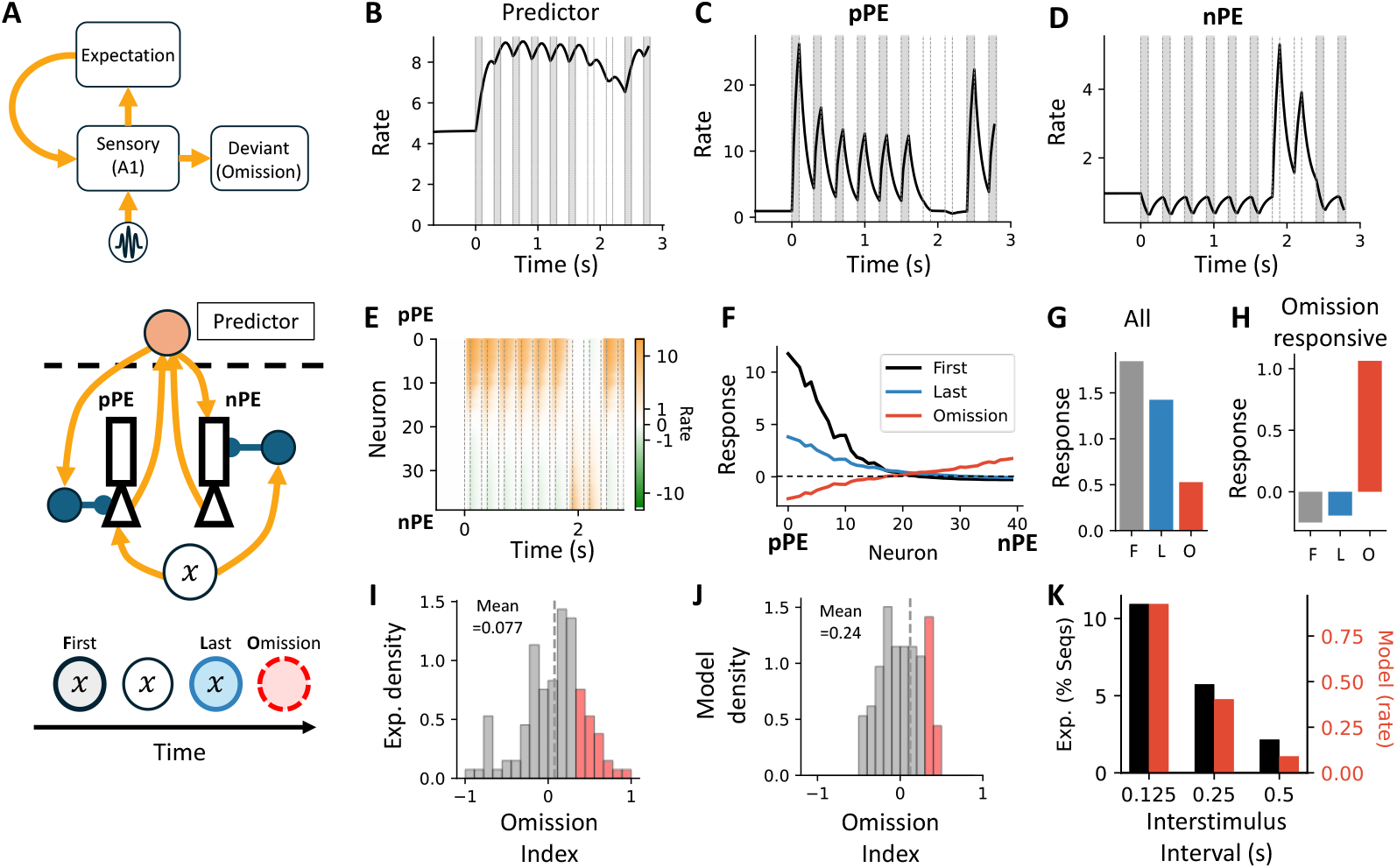
dPC reproduces omission responses, since the predictor input is net excitatory if presented in isolation. (A) Top: A1 can generate omission responses by internal expectation. Bottom: We use a predictor to generate the prediction input. The model receives several repetitions of the same stimulus to build up the prediction before the omission. (B) Predictor activity over time. The grey shading indicates the stimulus periods. (C, D) Time courses of a pPE neuron and an nPE neuron. The omission response is contributed by nPE neurons. (E) Response of individual neurons compared to the baseline. The neurons are sorted based on the strength of stimulus input. (F) Neural responses at the first occurrence, last repetition, and during omission. (G) Average response over the population. (H) Average response restricted to omission-responsive neurons. (I, J) Distribution of omission of data (I) and the model (J). (I) is modified from [27]. The dashed line indicates the mean. The red shading highlights neurons with omission index *>* 1*/*3, indicating their response is at least twice the response during the last repetition. (K) Population-level omission response as a function of the interstimulus interval. See [27] for the definition of omission-responsive sequence. (J, K) are generated in the realistic model version. Here, *τ_P_* = 0.1s.

In our model, the prediction input is net excitatory when presented in isolation, as in the omission case. This triggers the omission response naturally. In detail, the predictor starts with a high baseline firing rate of approximately 5 Hz [21] and saturates at 10 Hz. The predictor in simulation gradually builds up an internal expectation by integrating activity from the sensory area (Figure 2B), and generates corresponding top-down prediction input during stimulus periods. By our choice, a predictor with 10 Hz generates the maximum prediction input, as *P* = 1 in Figure 1E. During omission trials, predictor activity decreases but remains above baseline, driving a prediction signal. Though the net effect is excitatory for the whole population, it has differential effects on pPE and nPE neurons. During stimulus repetitions, pPE neuron activity decreases due to both stimulus-driven adaptation and top-down inhibition from the predictor (Figure 2C). During the omission period, their activity remains close to zero. In contrast, nPE neurons, excited by the predictor and inhibited by the stimulus, respond differently. During the omission, prediction input persists while stimulus-driven inhibition disappears, resulting in a strong excitatory response in nPE neurons (Figure 2D). At the population level, nPE neurons exhibit strong excited activity, while the low baseline firing rate limits suppression in pPE neurons (Figure 2E, F). Thus, on average, the overall population response to omission remains positive (Figure 2G). Among omission-responsive neurons (Figure 2H; see Methods), the response is higher than the population average during omission, but suppressed below baseline during the first and last repetitions, reflecting that the nPE neurons are inhibited by the stimulus.

In the experiment by [27], the authors quantified omission responses using an omission index, defined as the firing rate difference between the omission and repeated stimulus conditions, normalized by their sum. An omission index of 1 indicates that a neuron responds exclusively to the omission, whereas an index of –1 indicates exclusive response to the repeated stimulus. They found that neurons exhibited a broad distribution of omission indices with a positive mean (Figure 2I), and a subset of neurons had indices greater than 1*/*3, suggesting that their omission response was at least twice as large as their response to the repeated stimulus. We successfully reproduced these signature properties in the realistic model (see Methods, Figure 2J). In this version, each cell receives a different total maximum input, making the model more biologically plausible but less straightforward to interpret. We use the realistic model for comparisons with experimental data and explicitly indicate whenever it is applied. Additionally, [27] reported that doubling the inter-stimulus interval halved the proportion of neurons with significant omission responses, suggesting that the internal predictor may function as a leaky integrator. This behavior is reproduced in our realistic model and supports our modeling choice for predictor dynamics (Figure 2K).

### dPC reproduces key properties of mismatch negativity

Next, we study whether our dPC framework can explain the classic mismatch negativity (MMN) that is measured by the electroencephalogram (EEG) at the vertex of the scalp [48]. In a typical experiment, subjects are required to passively listen to a series of high-pitched tones and low-pitched tones and count the number of high-pitched tones in a block of 200 trials. The probability of high-pitch tones varied across blocks. When comparing the response to a high-pitched tone that follows a series of low-pitched tones to a control case in which the sequence contains only high-pitched tones, the EEG recordings show a larger response, reflected by the signature negative component peaking at 200 msec (N200), which is named as mismatch negativity (MMN). This signature is widely studied as a biomarker in diagnosing early psychosis [50]. Previous studies show that the MMN can be explained by a mass-type model [14, 15] that only describes the average population response and includes a top-down inhibitory effect through the pPE component. Does including the nPE component in dPC compromise the model’s ability to explain MMN?

The dPC framework claims that prediction input is net inhibitory when paired with a stimulus, reproducing the signature behaviors in MMN. In our model, we include *N_col_* = 8 columns, each tuned to a distinct stimulus frequency (Figure 3A), with a dedicated predictor. Each predictor receives excitatory input from *N_pCol_* = 40 neurons in its column and sends the prediction signal back. Within each column, we include both pPE and nPE neurons, responding to the corresponding stimulus and predictor. We begin with a case where a standard stimulus *x* appears with *p* = 0.7, and a deviant stimulus with *p* = 0.3. The predictors operate with a longer time constant *τ_p_* = 1 s, compared to the omission case. These predictors further inhibit each other via bell-shaped lateral inhibition (see Methods).

**Fig. 3.**
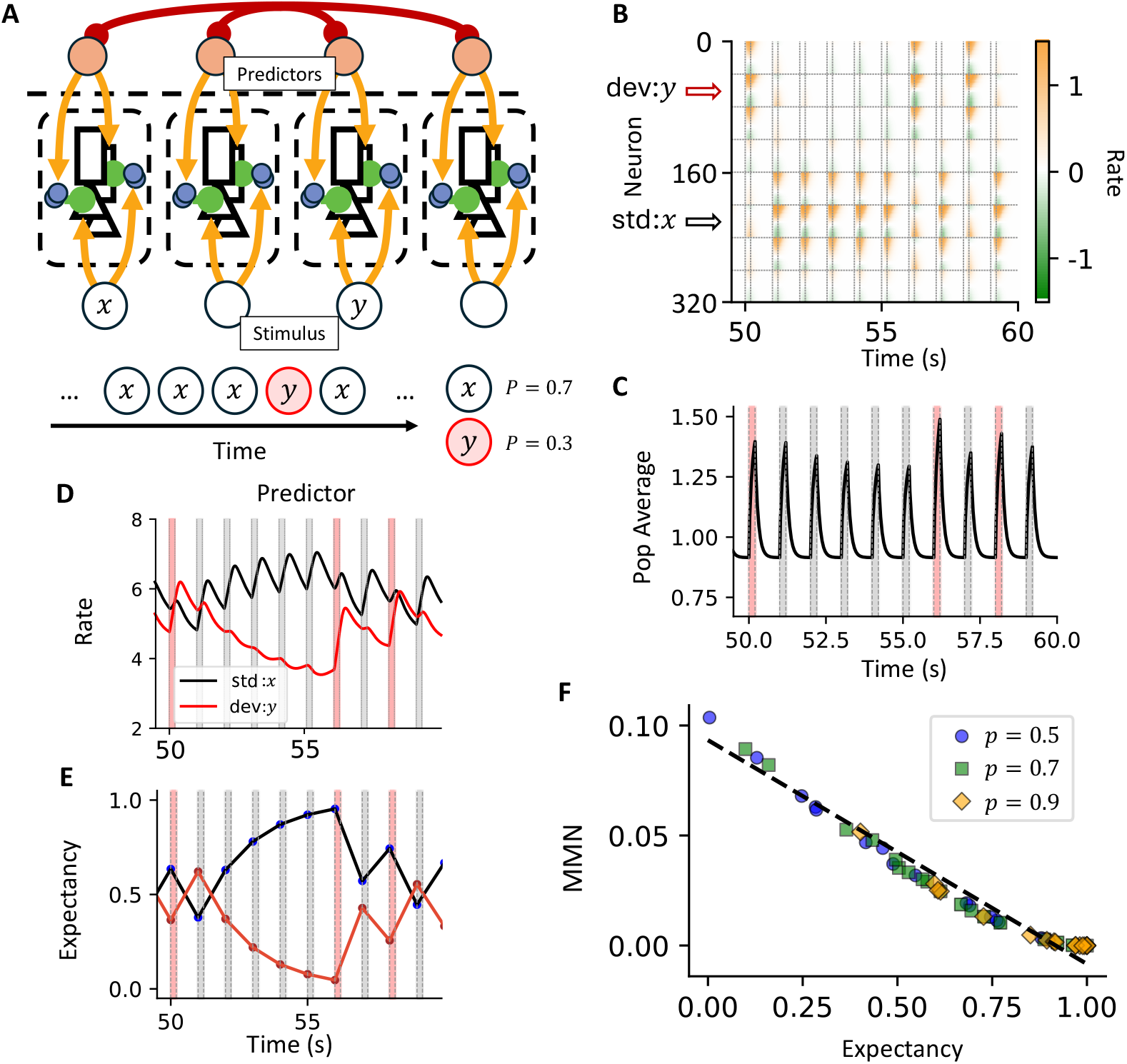
dPC reproduces key properties of mismatch negativity, since predictor input is net inhibitory if it co-exists with the stimulus. (A) Model schematic. Multiple columns of pPE and nPE neurons are included, each selective to different stimuli. Standard and deviant stimuli are presented randomly according to specified probabilities. (B) Firing rates of individual neurons over time. Horizontal dashed lines separate neurons by stimulus selectivity. Vertical dashed lines mark stimulus onset and offset. (C) Average firing rate across the population. Red shading indicates deviant stimuli; gray shading indicates standard stimuli. (D) Predictor activity for both standard and deviant stimuli. (E) Expectancy of the standard and deviant stimuli over time. See Methods. (F) MMN is negatively correlated with expectancy. See the main texts for details. Different symbols represent data from different simulations. Symbols are downsampled for clarity, but fitting is performed on all data combined across runs. Here, *τ_P_* = 1*s*.

In this model, individual neurons respond heterogeneously to the stimulus (Figure 3B). Neurons from different columns are stacked vertically, with each column separated by horizontal dashed lines. Within each column, pPE neurons have smaller indices than nPE neurons. As expected, pPE neurons in the stimulus-matched column are strongly activated during the stimulus period, while nPE neurons within the same column are slightly suppressed. At the population level, the average activity in each column remains positive, as pPE excitation outweighs nPE suppression. In addition, the average population response varies with stimulus history (Figure 3C). Activity is highest for the deviant stimulus *y*, while responses to repeated presentations of the standard stimulus *x* gradually decline. This modulation reflects changes in predictor activity over time (Figure 3D): as *x* is repeatedly presented, predictor*_x_*activity builds up, leading to reduced activity in column *x*, since the prediction input is net inhibitory when paired with its corresponding stimulus. This pattern reverses when a deviant stimulus *y* is presented. Due to lateral inhibition, predictor*_y_*activity remains low while predictor*_x_*stays elevated. The resulting reduced inhibition from predictor*_y_*leads to a stronger response in column *y*. Notably, column *x* also shows a positive response when the deviant stimulus *y* is presented, as predictor*_x_*excites column *x* in the absence of stimulus *x*, mimicking an omission response. This will be discussed further in later sections.

A previous study [48] explained the MMN in a Bayesian framework by a latent “expectancy” variable, which estimates leaky stimulus history using a fitted memory decay rate *α* (Supplementary Figure 9A, see Methods). To test whether this expectancy also accounts for the activity changes in the model, we quantified MMN as the difference in population responses to a stimulus presented in a randomized sequence versus a fixed sequence containing only that stimulus (i.e., *p* = 1). We found that MMN is linearly and negatively correlated with the latent expectancy (Figure 3E, F), agreeing with the experimental observation (Supplementary Figure 9A).

The degree of leakiness in calculating the expectancy is controlled by the decay rate *α*. To assess its robustness, we evaluated model fit using the explained variance *R*^2^ (Supplementary Figure 9B). We found that *α* ∈ [0.5, 0.7] consistently produces a strong anti-correlation between expectancy and MMN. With our carefully chosen parameters, the optimal decay rate is *α*_leak_ = 0.6, which matches the value reported in experimental data [48] (Supplementary Figure 9A).

To better illustrate how MMN increases with repetition, we tested our model using a fixed sequence of *N_rep_* standard stimuli followed by a single deviant (Supplementary Figure 4). The longer the repetition sequence, the less expected the deviant stimulus. Here, stimulus presentation triggered heterogeneous responses across individual neurons (Supplementary Figure 4A, D), as seen previously. When comparing *N_rep_* = 6 to *N_rep_* = 2, the activity difference is larger across predictors dedicated to the standard and the deviant stimulus (Supplementary Figure 4B, E), resulting in a stronger MMN (Supplementary Figure 4C, F). MMN amplitude saturates around *N_rep_* = 5 for predictor time constant *τ_P_* = 1 (Supplementary Figure 4G, as used in Figure 3). A smaller predictor time constant leads to faster MMN saturation with fewer repetitions (Supplementary Figure 4G), but the MMN saturates at a smaller amplitude (Supplementary Figure 4H).

### dPC predicts nPE contributions in deviance detection tasks

Though MMN is often explained through predictive coding, an alternative account attributes it to fatigue-like mechanisms, such as stimulus-specific adaptation (SSA) [15]. To isolate top-down contributions, recent studies have focused on deviance detection (DD), which measures context modulation by comparing responses to the same stimulus presented either as a deviant or as a probability-matched control [16, 17, 5]. The control is typically the first occurrence of the stimulus (Figure 5A) or a version interleaved within a sequence of randomized stimuli, both minimizing adaptation effects. In this section, we focus on using the first occurrence as the control, and address the randomized control in the next section. In addition to the population activity, advances in techniques like calcium imaging now enable measurement of individual neuron activity. While deviance detection at the single-neuron level is variable and heterogeneous [16, 17], can this heterogeneity be dismissed as random noise, as in mass models [14] that describe only population-averaged responses? Or is it functionally meaningful for computing predictive error?

**Fig. 4.**
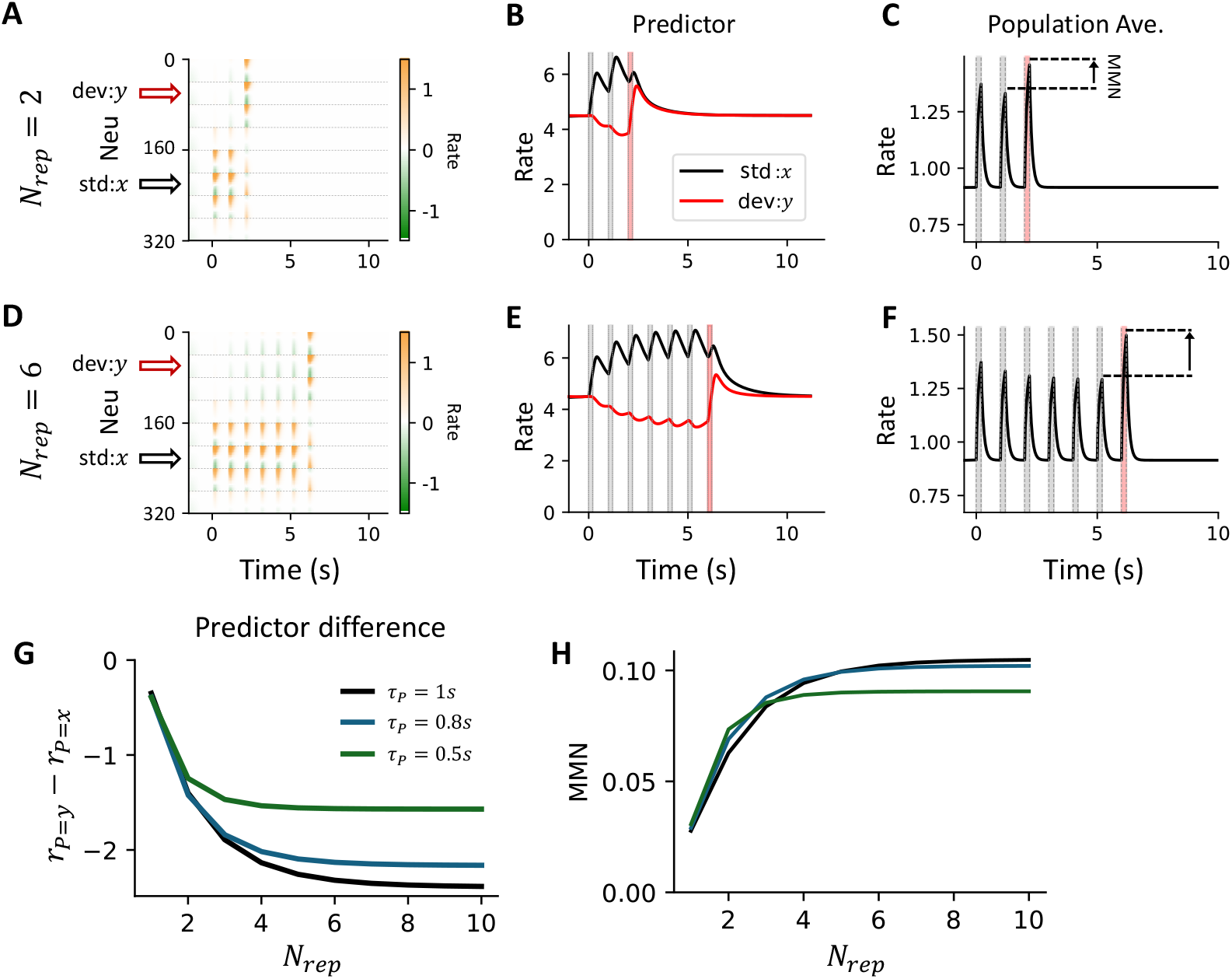
MMN increases with the number of repetitions *N_rep_*. Related to figure 3. (A) Activity of individual neurons with two repetitions of the standard stimulus (*N_rep_*= 2). (B) Predictor activity for both standard and deviant stimuli. (C) Average population activity over time. (D to F) Same as (A to C) but with six repetitions (*N_rep_*= 6). (G) The difference in predictor activity between deviant and standard stimuli (*r_P_* _=*y*_ − *r*_*P*=*x*_) becomes more negative as the number of repetitions increases. Each color represents a different *τ_P_* value. A longer time constant leads to slower saturation of the predictor signal. (H) MMN increases over the number of repetitions.

**Fig. 5.**
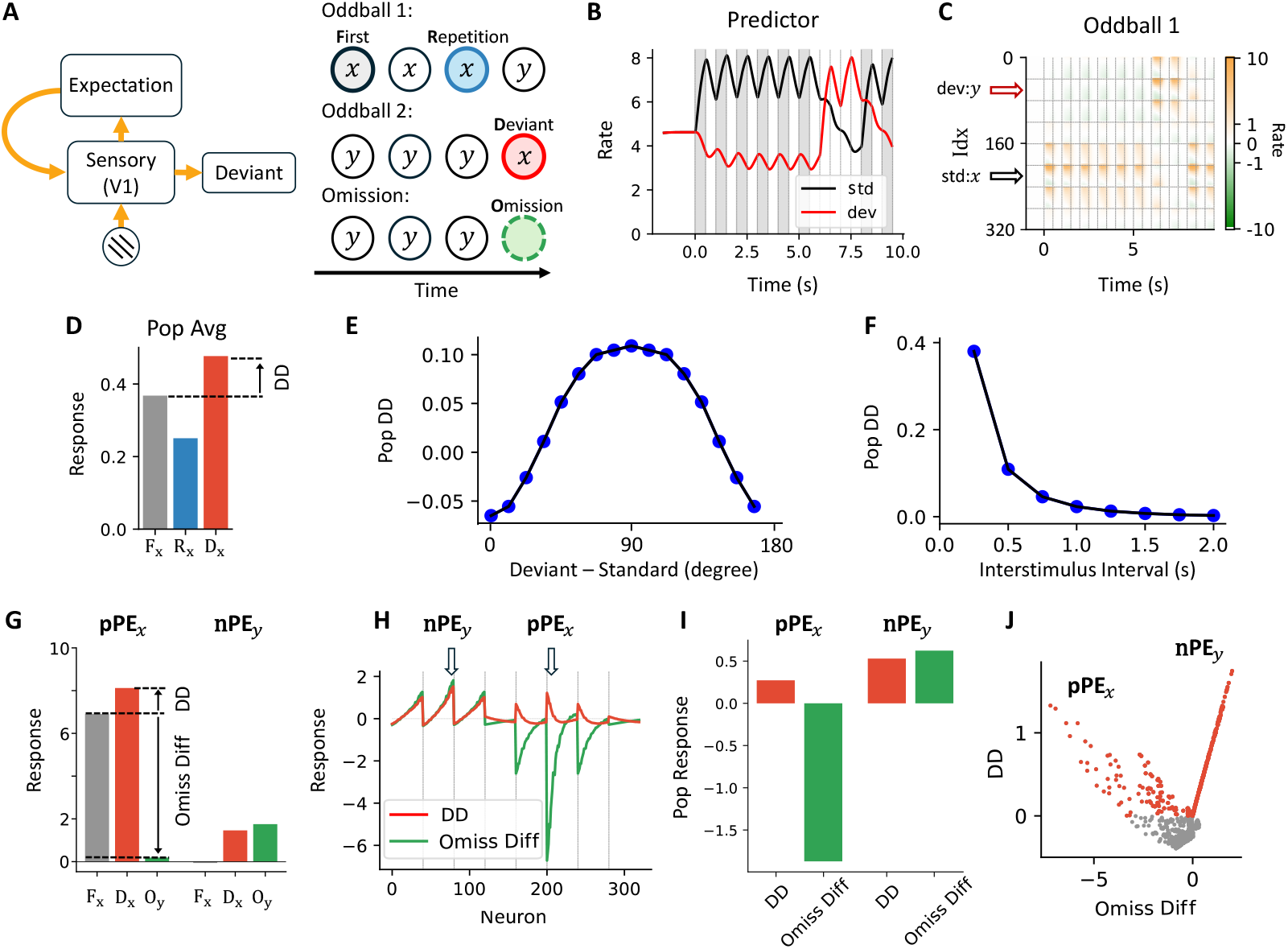
dPC predicts functional role of nPE neurons in deviant detection. (A) Left: V1 can generate deviance detection signals in an oddball protocol. Right: simulation schematic. Responses to the same stimulus presented in different contexts (first, repetition, deviant, or omission) are compared. (B) Predictor activity in the corresponding standard (black) and deviant sequences (red). (C) Firing rates of individual neurons in oddball sequence 1. (D) Average population activity across different contexts within the oddball sequences. (E) Population DD response decreases as the orientation of the deviant stimulus diverges from the perpendicular phase. (F) The population DD response decreases with increasing stimulus interval. (G) Firing rates of an example pPE*_x_* neuron and an example nPE*_y_*neuron when stimulus *x* is presented as the first occurrence, as a deviant, or is omitted. (H) Single neuron responses. The red line indicates the deviance detection, and the green line indicates the omission difference, calculated as the difference between the omission response and the response at the first occurrence. (I) DD and omission differences of pPE*_x_* and nPE*_y_*population. While both show positive DD responses, only the nPE*_y_*population exhibits an omission response greater than the first occurrence. Scatter plot of DD versus omission difference. (I, J) are generated from the realistic model version. See Methods. Here, *τ_P_* = 0.4*s*.

Our dPC framework reproduces DD responses at both the population and single-neuron levels in sensory areas such as V1. The network structure remains the same as in Figure 3A, but we use a shorter predictor timescale (*τ_P_* = 0.4 s) and include stimulus adaptation across repetitions (see Methods). To test whether neural responses reflect contextual change, we simulate two oddball sequences: sequence no. 1 (*xxxy*) where stimulus *x* appears repeatedly until the deviant *y*, and sequence no. 2 (*yyyx*) where *x* is the deviant (Figure 5A). Stimuli *x* and *y* represent orthogonal grating orientations. We also test omission responses using the sequence *yyyo* with the same number of repetitions in oddball sequence no.1 and no.2.

In later sections, we label stimulus presentations in different contexts as follows: F*_x_* denotes the first occurrence of stimulus *x*, R*_x_* denotes the fourth or later repetitions, and D*_x_* indicates that stimulus *x* is presented as a deviant. Analogous notations F*_y_*, R*_y_*, and D*_y_* apply to stimulus *y*. Omissions in sequences *xxxo* and *yyyo* are labeled O*_x_* and O*_y_*, respectively. We refer to oddball sequence no. 1 as the standard sequence for *x* and oddball sequence no. 2 as the deviant sequence, with terminology applied symmetrically for *y*.

As shown earlier in the MMN case, predictor activity increases across repetitions when the corresponding stimulus is presented in the standard sequence, but decreases when the corresponding stimulus is presented in the deviant sequence due to lateral inhibition (Figure 5B). Their activities, which are different across contexts, modulated the neuron response to the same stimulus (Figure 5C). As a whole population, the sensory response to R*_x_* is below that to F*_x_*, and it is larger to D*_x_* than to F*_x_* (Figure 5D). Population-level deviance detection to stimulus *x* (DD), quantified as the response difference to D*_x_* and F*_x_*, varies with the angular difference between stimuli *x* and *y* (Figure 5E). Maximum deviance detection occurs when *x* and *y* are orthogonal. As the angular difference decreases, DD diminishes. When the angular difference reaches zero, the DD flips sign, reflecting that D*_x_* and R*_x_* become the same. Similar to the trend observed in the omission protocol, DD decreases exponentially with increasing inter-stimulus interval, reflecting the decaying memory of the predictor (Figure 5F). The prediction on how DD changes over the angular difference and interstimulus interval can be tested in future experiments.

At the single-cell level, unlike the context-dependent modulation seen at the population level, the predictor consistently inhibits pPE neurons and excites nPE neurons within the same column (Supp. Figure 10A, B). Since predictor activity is higher during R*_x_* than F*_x_*, and lower during D*_x_* than F*_x_*, the pPE*_x_*neuron is more inhibited in the repeated condition and less inhibited in the deviant condition.

In contrast, the nPE*_x_* neuron is more excited during R*_x_* and less excited during D*_x_*, though its overall response is smaller than that of pPE*_x_*. The activity of all neurons in the *x*-selective column is shown in Supp. Figure 10C, D. Since the reduced response to R*_x_* could arise from both stimulus-specific adaptation and top-down modulation, we tested a condition without adaptation (Supp. Figure 10E). In this case, the reduction in response to R*_x_* is attenuated, while the increase in response to D*_x_* remains unchanged, suggesting that DD may be a more reliable readout of top-down prediction than MMN.

As suggested in the previous section, the population-level DD response during D*_x_* arises from two distinct sources in the model. The first is reduced inhibition from the predictor in the *x*-selective column, as previously discussed. The second originates from the omission-like response in the *y*-selective column, where predictor*_y_*exerts a net excitatory effect, particularly on nPE*_y_*. Notably, nPE*_y_* shows a DD response comparable to pPE*_x_*, but its response to F*_x_* is much weaker (Figure 5G). This highlights a potential confound: if we define DD solely by comparing D*_x_* and F*_x_*, we might mistakenly classify nPE*_y_* as a pPE*_x_*neuron, even though nPE*_y_* does not respond to stimulus *x*.

Next, we find that a commonly utilized omission protocol can evoke responses that clearly differen-tiate pPE*_x_*and nPE*_y_* neurons (Figure 5A, G), as only nPE*_y_* neurons respond to omission O*_y_*, whereas pPE*_x_*neurons do not (Figure 5H). In simulations using the realistic model, both populations exhibit DD responses, but only nPE*_y_* neurons respond more strongly to O*_y_* than to F*_x_*. When plotting individual neuron activity along DD magnitude and response to O*_y_*, the two populations form distinct branches (Figure 5J). This separation between pPE*_x_*and nPE*_y_* provides an experimental prediction that should be tested in future studies.

Lastly, a detailed decomposition of input contributions to both pPE*_x_*and nPE*_y_* across different protocols is shown in Supplementary Figure 11.

### dPC predicts distinct pPE and nPE responses in a randomized oddball protocol

The difference between pPE*_x_*and nPE*_y_* neurons can be further tested in a randomized oddball protocol by comparing responses to the first occurrence (F*_x_*) and to a control condition (C*_x_*) in addition to using the omission protocol (Figure 6A). Here, the first occurrence F*_x_* refers to the presentation of standard stimulus *x* immediately following the deviant stimulus *y* within a long sequence, not the first appearance of *x* in the entire run (Figure 6A). In [16, 17, 5], a many-standard sequence was used as the control instead of the first occurrence to avoid potential novelty effects. In the many-standard control, eight differently oriented gratings, including those used in the oddball sequences, are presented with equal probability. This design helps eliminate the influence of adaptation and expectation signals on neural responses. The oddball sequences are randomized such that the number of repetitions between two deviants varies, rather than being fixed, to further minimize expectation-related effects. Here, we apply the same protocol to test our model. Individual neuron responses in the many-standard control sequence are shown in Figure 6B.

**Fig. 6.**
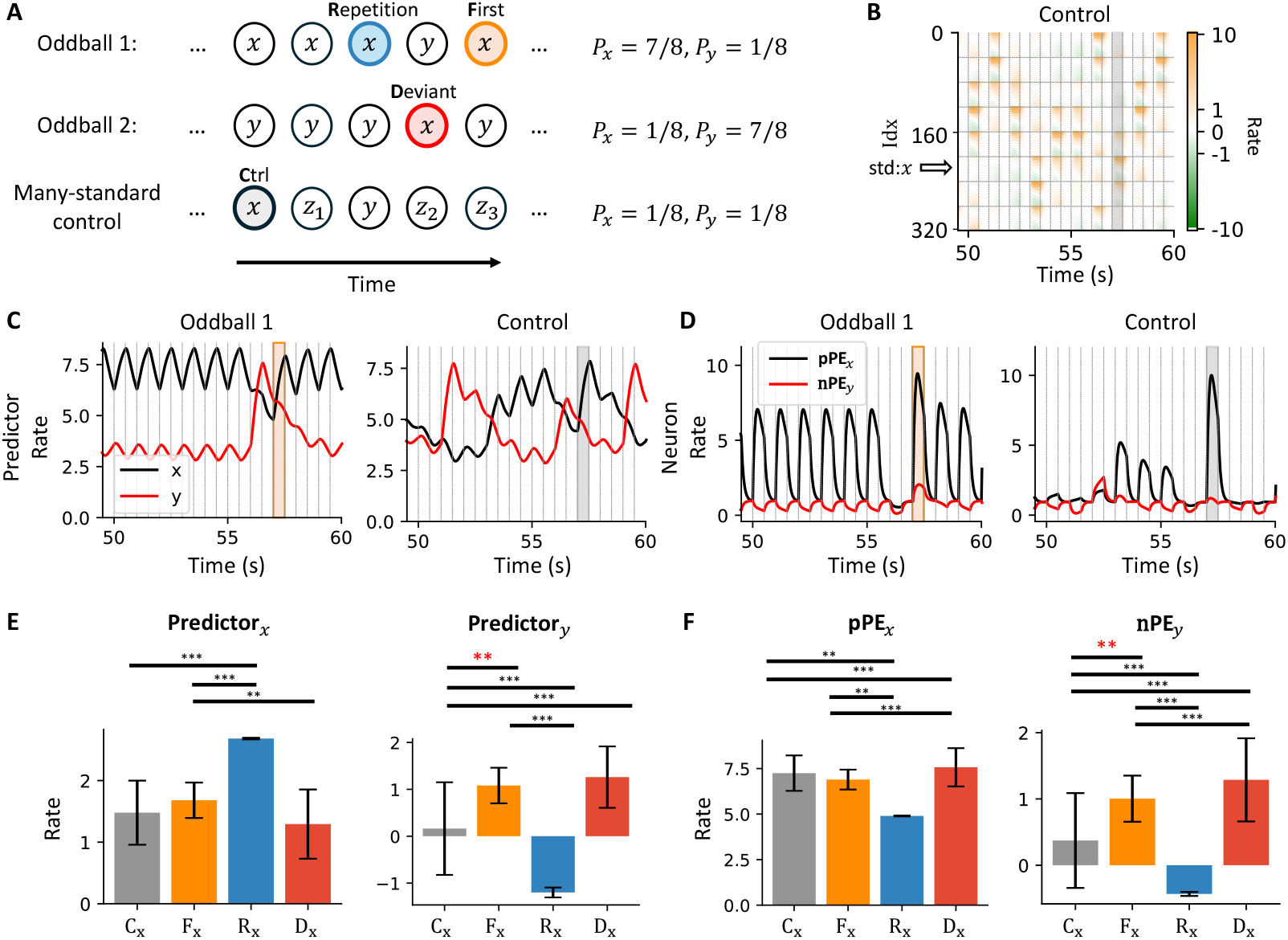
nPE*_y_*neurons show stronger responses to F*_x_*than it to C*_x_*. (A) Simulation schematic. The comparison focuses on the first stimulus presentation versus the stimulus in the many-standard control condition. (B) Activity of individual neurons in the many-standard control condition. The gray shading indicates when stimulus *x* is presented. (C) Predictor activity during oddball sequence one (left) and the many-standard control (right). Orange shading indicates the F*_x_*, and gray shading indicates the C*_x_*. Activity for predictor*_x_* and predictor*_y_* is shown. (D) One pPE*_x_* and one nPE*_y_*activity during the oddball sequence 1 (left) and many standard controls (right). (E) predictor*_x_*and predictor*_y_*responses compared to the baseline across the C*_x_*, F*_x_*, R*_x_*and D*_x_*contexts. Activity of predictor*_y_* is significantly more larger during F*_x_*than during C*_x_*. The error bar indicates the standard deviation, which indicates the variance of the response under the same context. The standard error of the mean (SEM) is essentially zero due to the large sample size of stimulated data. ^∗∗^: *p <* 0.01; *^∗∗∗^*: *p <* 0.001. (F) Activity of a pPE*_x_* and nPE*_y_*neuron across conditions. The activity of nPE*_y_*neuron is significantly larger during F*_x_*than during C*_x_*. Detailed *p*-values are provided in the Supplementary Materials.

**Fig. 7.**
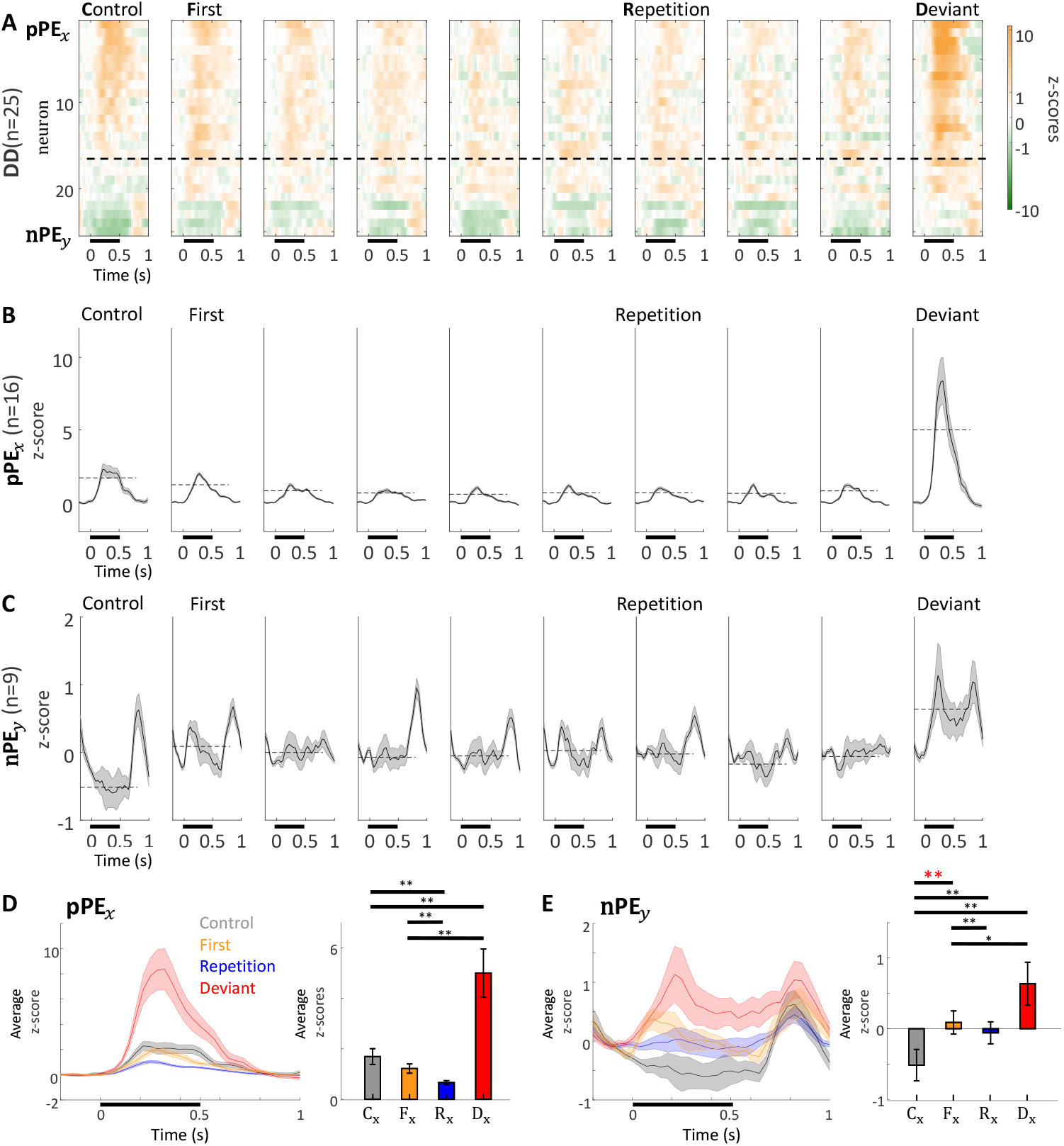
Putative pPE*_x_* and nPE*_y_*neurons are both observed in an oddball paradigm experiment. (A) Activity of all identified deviant detector (DD) neurons (*n* = 25) in the recorded population. DD neurons are defined as those that respond significantly more strongly to the deviant than to the control stimulus (*p <* 0.05). Neurons above the dashed line also show a significant response in the many-standard control condition (*p <* 0.05) and are considered as pPE*_x_*neurons. The remaining neurons, which do not respond significantly in the control, are classified as nPE*_y_*neurons. Neurons are sorted based on average activity during the control period. The black bar below the x-axis indicates the stimulation period. (B) Average z-scored activity of pPE*_x_*neurons (*n* = 15) across different contexts. The shading indicates the SEM over time. (C) Average z-scored activity of nPE*_y_*neurons (*n* = 9) across different contexts. (D) (Left) Overlay of z-scored pPE*_x_*neuron responses across contexts. (Right) Mean z-score during the stimulus period. The error bar indicates the SEM. No significant difference is observed between C*_x_*and F*_x_*(*p >* 0.05). (E) Same as (D), but for nPE neurons. The activity during F*_x_*is significantly larger than during C*_x_. ^∗^*: *p <* 0.05; ^∗∗^: *p <* 0.01; *^∗∗∗^*: *p <* 0.001.

**Fig. 8.**
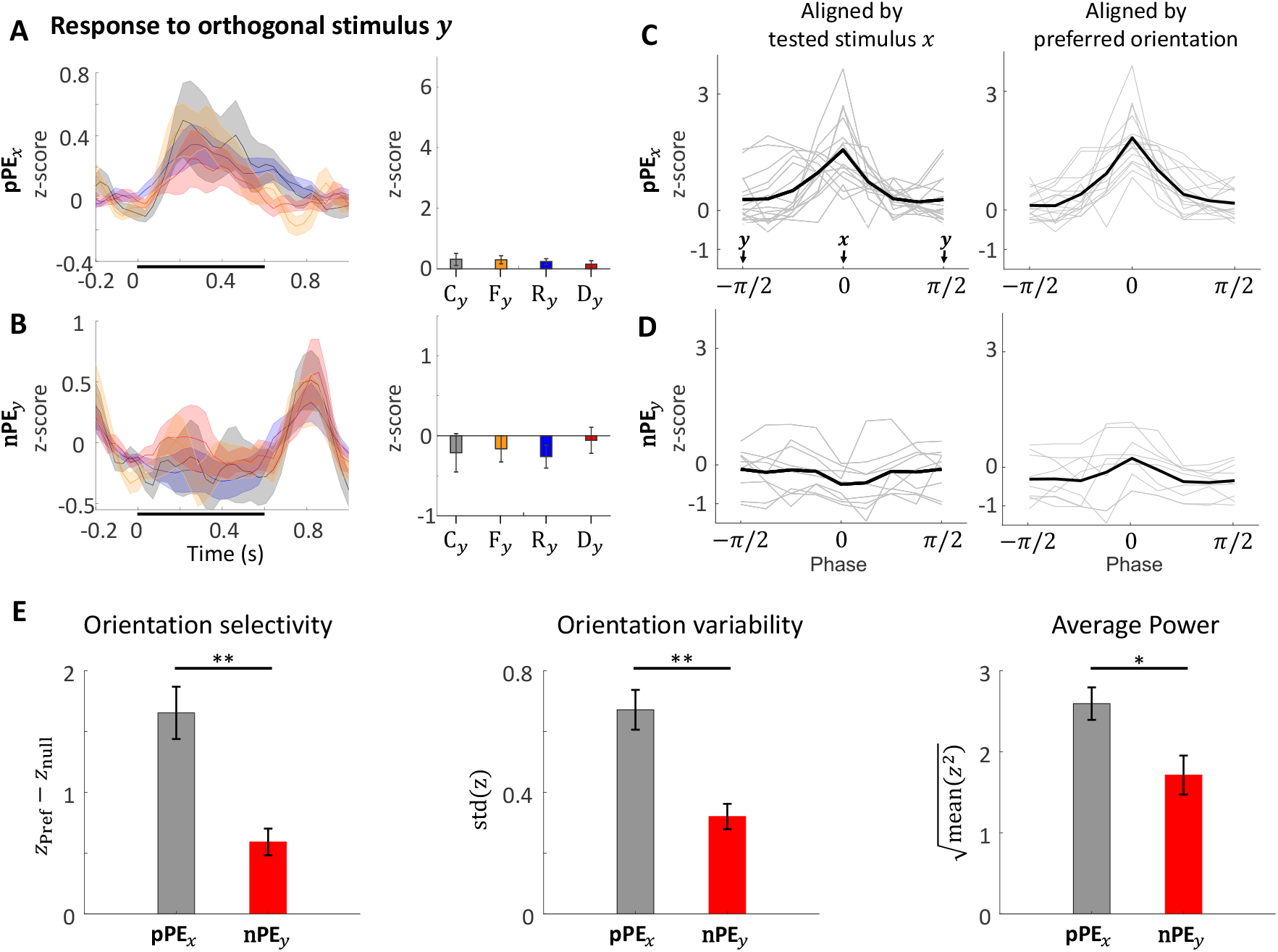
Selectivity of pPE*_x_* and nPE*_y_*neurons. (A) Overlay of z-scored pPE*_x_* neuron responses to stimulus *y* across contexts (Left). See Supplementary Figure 12G, H for responses in different contexts separately. (Right) Mean z-score during the stimulus period. The error bar indicates the SEM. No significant difference is observed between any pair of responses. (B) Same as (A) but for nPE*_y_*neurons. (C) Average z-scored responses to different stimulus orientations for pPE*_x_*neurons. Gray lines represent tuning curves for individual neurons; the black line shows the population average. (Left) The tuning curves are aligned with the tested stimulus *x* in the oddball protocols. Stimulus *y* is orthogonal to stimulus *x*. Stimulus *x* and *y* are marked with arrows. Stimulus *y* shows twice to show periodicity. (Right) The same tuning curves aligned with the preferred orientation of each neuron. (B) Same as (A), but for nPE*_y_*neurons. (E) Orientation selectivity (left), Orientation variability (middle), and average power (right) forpPE*_x_* and nPE*_y_*neurons. Orientation selectivity index is calculated as the z-score at the preferred orientation minus the z-score at the orthogonal (null) orientation. Error bars represent the SEM. Note that the preferred orientation is not necessarily at 0 degree. ^∗∗^: *p <* 0.01. Orientation variability is measured as the standard deviation of z-scores across all tested orientations. Average response power is computed as the geometric mean of z-scores across orientations. *^∗^*: *p <* 0.05. Detailed *p*-values are reported in the Supplementary Materials.

As before, predictor activity reflects a leaky integration of stimulus history (Figure 6C). In oddball sequence no. 1, since stimulus *x* occurs more frequently, predictor*_x_*maintains elevated activity throughout most of the sequence, while predictor*_y_*becomes active only when stimulus *y* appears. By definition, F*_x_* follows D*_y_*. During F*_x_*, predictor*_x_*begins to ramp up, and predictor*_y_*starts to decline.

However, due to the predictors’ long timescale, predictor*_y_* remains elevated during F*_x_* compared to R*_x_*. In contrast, under the many-standard control protocol, predictor activity stays near baseline unless the corresponding stimulus is presented, reflecting the randomized nature of the sequence.

When comparing predictor activity across all trials and contexts, the predictor*_x_*does not differ significantly between C*_x_* and F*_x_*, whereas predictor*_x_*shows a significant increase during F*_x_* (*p <* 0.01, Welch’s t-test; Figure 6E). This pattern is mirrored in the responses of pPE*_x_*and nPE*_y_* neurons: only the nPE*_y_* neuron shows a significantly higher response (*p <* 0.01) during F*_x_* than C*_x_*, but not the pPE*_x_*neuron (Figure 6D, F).

All pairwise *p*-values are reported in the Supplementary Materials.

**Fig. 9.**
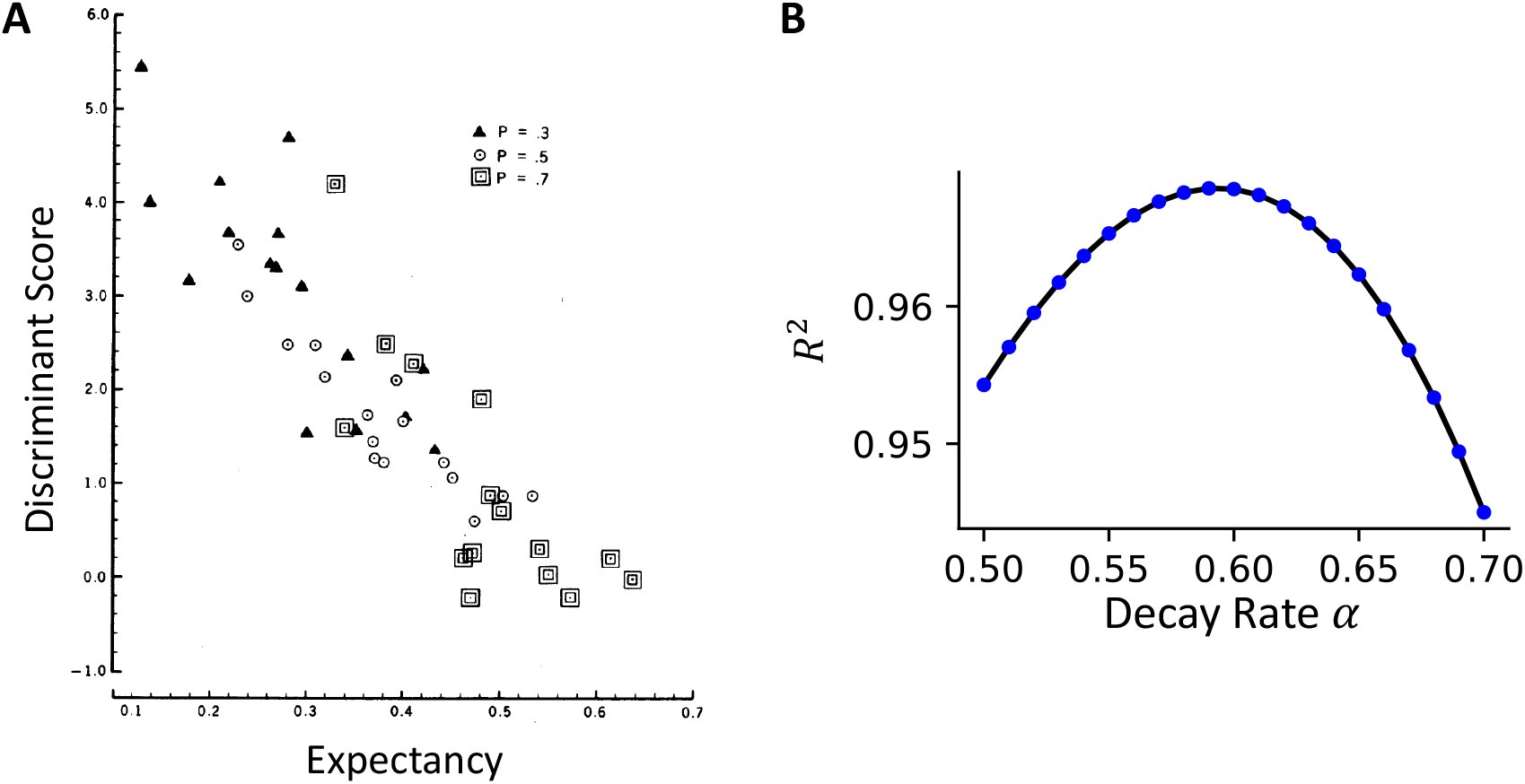
Related to figure 3. (A) Model behavior is best captured with a memory decay rate of *α* = 0.6. The y-axis shows the variance explained by a linear fit between the difference in population-averaged activity and the computed expectancy. (B) Relationship between discriminant score and expectancy, reproduced from [48] with permission. The discriminant score is calculated based on the N200, P300, and slow waveform components. Expectancy is computed using a memory decay rate of *α* = 0.6.

**Fig. 10.**
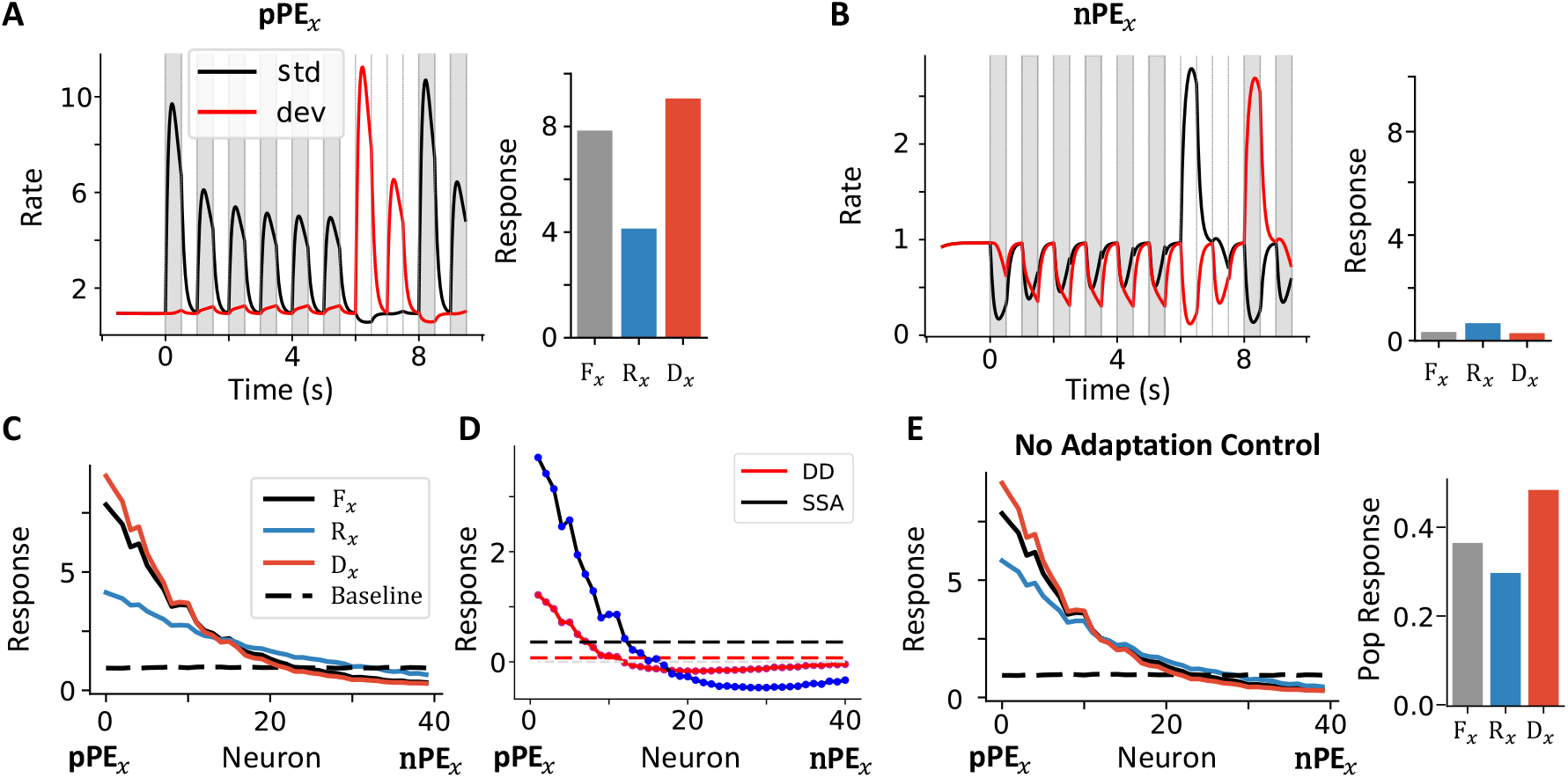
Individual neuron activity in fixed oddball sequence and no-adaptation control. Related to figure 5. (A) a pPE*_x_*neuron over time when the corresponding stimulus is shown as the standard (black) or deviant stimulus (red). (B) an nPE*_x_*neuron activity. (C) Population response across different contexts. (D) deviance detection (DD, deviant minus first occurrence) and stimulus-specific adaptation (SSA, first minus last repetition) of all the neurons. Dashed lines indicates corresponding average response of the population. (E) No adaptation control. Here, the stimulus is not adapted. See Methods. The individual (left) and averaged population response across all columns (right). Compared to the case with stimulus adaptation, only the activity during the last repetition increases, while the deviant response remains unaffected.

**Fig. 11.**
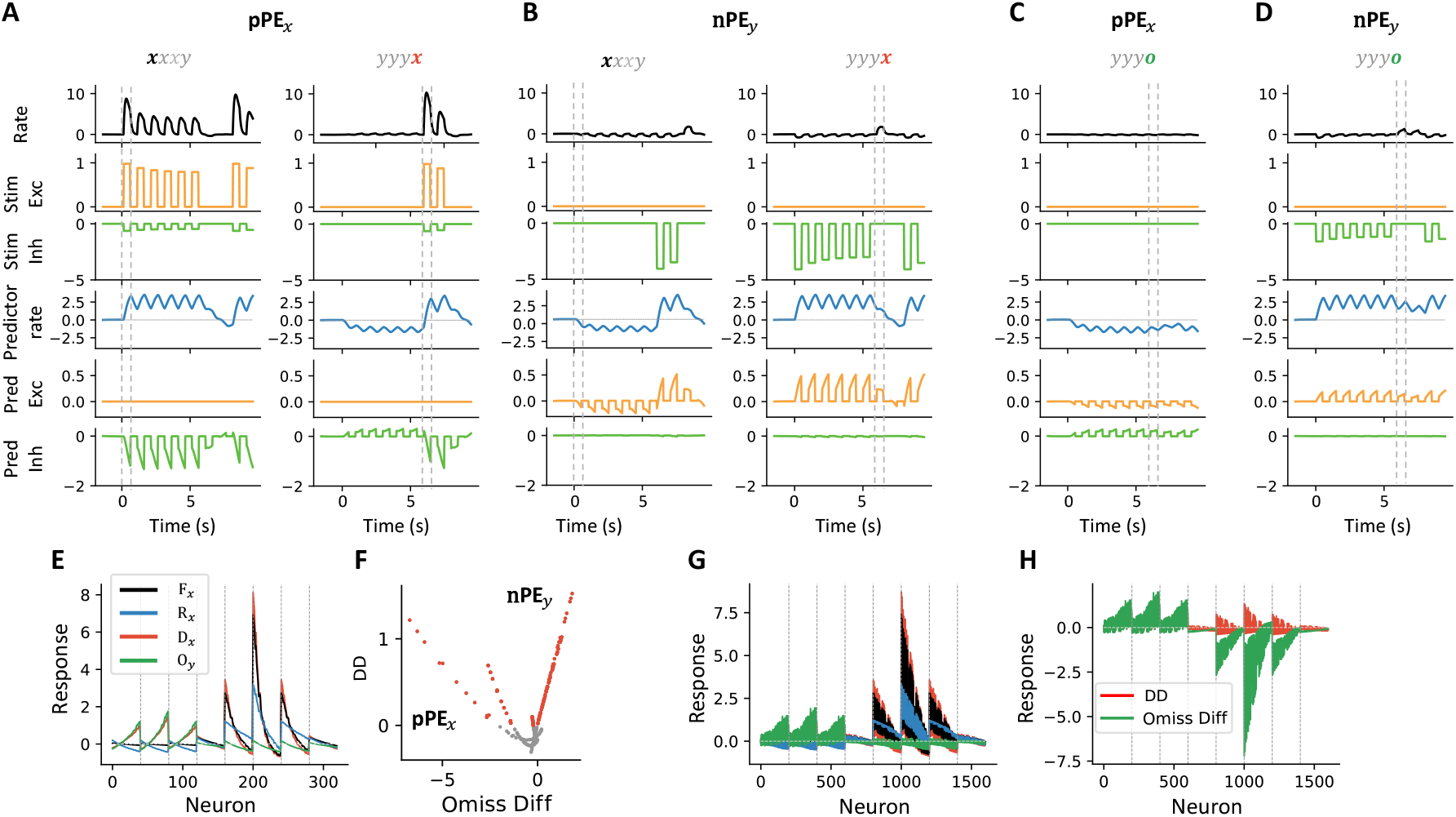
Input differences across channels between pPE*_x_* and nPE*_y_*neurons. Related to figure 5. (A) Activity of pPE*_x_*during the fixed oddball sequences *xxxy* and *yyyx*. From top to bottom: neuron firing rate; somatic excitation, representing stimulus-driven excitation; dendritic inhibition relayed by the I2 population, representing stimulus-driven inhibition; predictor activity selective to stimulus *x*; dendritic excitation, representing prediction-related excitation; and somatic inhibition relayed by the I1 population, representing prediction-related inhibition. (B) Same as (A), but for a nPE*_y_*. (C, D) Same layout as (A), but in the omission protocol *yyyo*. (E) Individual neuron responses across contexts. (F) Scatter plot of deviance detection (DD) and omission difference in the simplified model version, which assumes the total maximum input is the same across neurons. (G) Same as (E) but based on results from the realistic model version. (H) Deviance detection and omission difference in the realistic model version (Related to Figure 5J).

**Fig. 12.**
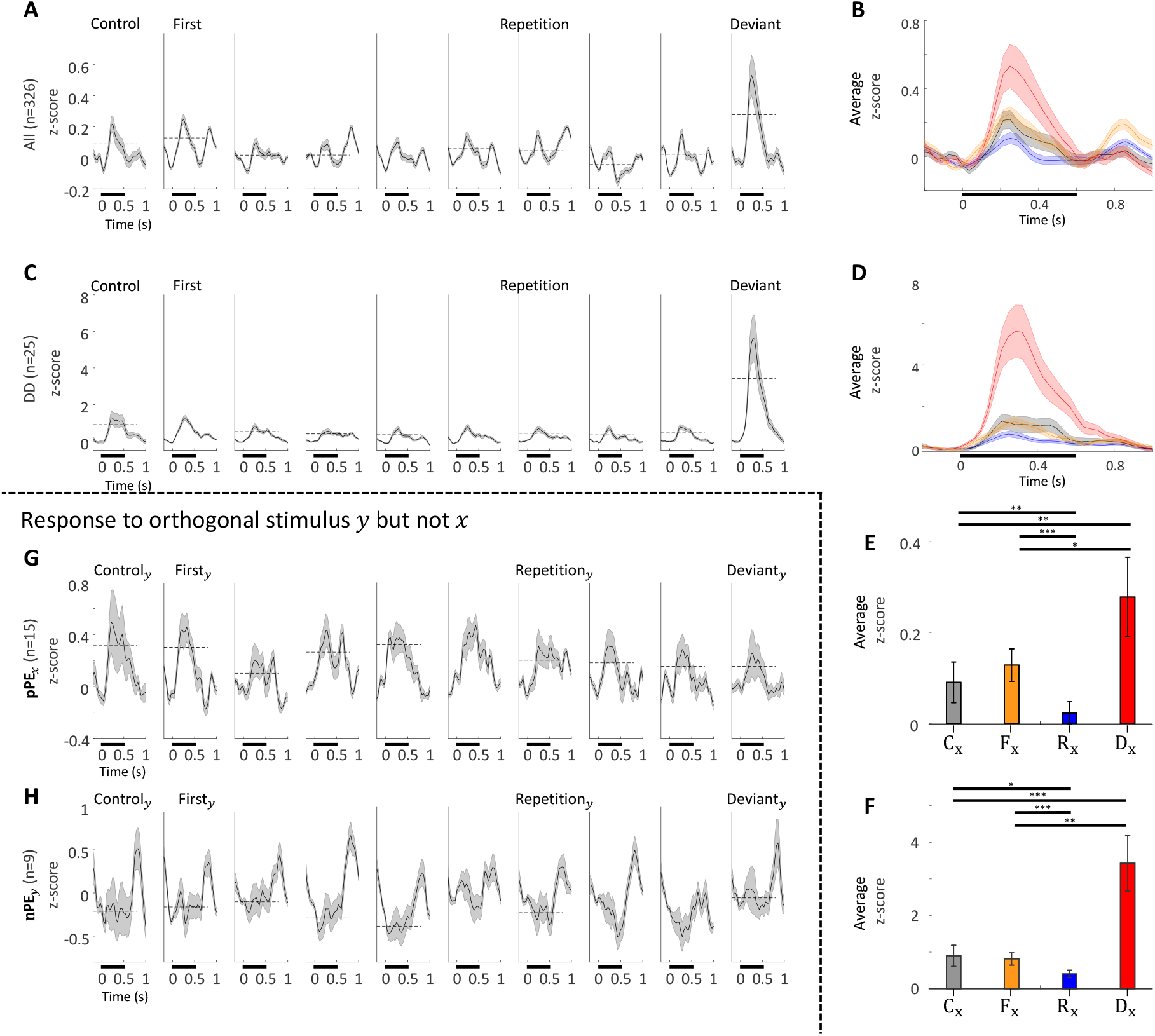
Further analysis on neurons during randomized oddball protocols. Related to Figure 7. (A) Average z-scored activity of all recorded neurons across different contexts. The black bars below the x-axis indicate the stimulation period. (B) Overlay of neural activity traces across contexts. (C, D) Same as (A, B), but limited to deviant detectors (*n* = 25). (E, F) Mean activity of the whole population and DDs across contexts, consistent with the observation in [5]. *^∗^*: *p <* 0.05; ^∗∗^: *p <* 0.01; *^∗∗∗^*: *p <* 0.001. (G) pPE*_x_*responses to stimulus *y*, a grating that is perpendicular to the stimulus *x* in the oddball protocol by reversing the sequence no.1 and no.2. No significant differences are observed across contexts. (H) Same as (G), but for nPE*_y_*neurons. Again, no significant differences are found across contexts. Detailed *p*-values are reported in the Supplementary Materials.

### dPC predictions are confirmed in the experiment

Our model pinpoints predictions that can be experimentally tested. We have argued that both pPE*_x_*and nPE*_y_* neurons contribute to deviance detection and, therefore, should both exhibit stronger responses to deviant stimuli than to the many-standard control. Additionally, in contrast to pPE*_x_*neurons, nPE*_y_*, responding to [P*_y_ −* S*_y_*]_+_, should not respond to the tested stimulus in the control condition C*_x_*. Furthermore, nPE*_y_* neurons should respond more strongly during F*_x_* than during C*_x_*.

We tested these predictions using both previously published data [5] and newly collected recordings, comprising a total of *N* = 326 stimulus–neuron pairs from three mice of either sex. The average responses of these neurons are shown in Supplementary Figure 12A, B, E. In these experiments, a 0.5s stimulus was presented to the animals with jittered inter-stimulus intervals ranging from 0.4 to 0.6s. The detailed experimental procedure is described in [5].

Among these neurons, 25 out of a total *N* = 326 showed significantly stronger responses in the deviant context compared to the control (*p* < 0.05, Welch’s t-test; Figure 7A, Supplementary Figure 12C, D, F). We refer to these as deviant detectors (DDs). This proportion is consistent with other reports during basic oddball paradigms [17, 5]. Within the DD population, 16 neurons also showed significant responses to the control condition (*p <* 0.05, paired t-test), while the remaining 9 did not (*p >* 0.05). We classify the former as pPE*_x_*neurons and the latter as nPE*_y_* neurons, since only pPE*_x_*neurons are expected to respond in the control condition.

On average, the pPE*_x_*neurons reproduce the response pattern previously reported [16, 17, 5] (Figure 7B), showing similar responses during C*_x_* and F*_x_*. Their activity decreases with repeated presentations but increases in D*_x_*. In contrast, the nPE*_y_* neurons (Figure 7C) exhibit a slightly suppressed response in the control condition. Their responses are larger during F*_x_* and even more elevated during D*_x_*.

When examining average responses across the stimulus period (Figure 7D), pPE*_x_*neurons do not show a significant difference between C*_x_* and F*_x_*. In contrast, nPE neurons have significantly higher activity during F*_x_* than C*_x_* (*p* < 0.05, paired t-test; Figure 7E), reflecting that nPE neurons are excited by the top-down prediction input.

In addition, we observed several differences in the average response time course. For nPE*_y_* neurons, response onsets in the deviant context often occur slightly (about 100ms, or 3 frames in the recordings) before stimulus onset (Figure 7E), which is consistent with the idea that prediction signals arrive in advance of sensory input, reflecting internal expectation. These neurons also exhibit a rebound in activity following the stimulus period. One possible explanation is that the nPE*_y_* population is nonspecifically inhibited by the stimulus (i.e., a nPE*_y_* neuron can be inhibited by both stimulus *x* and *y*), and that removal of the stimulus disinhibits the neurons, leading to rebound activity.

### pPE and nPE are further separated through selectivity

pPE*_x_*and nPE*_y_* neurons are further separated based on their responses to gratings of different orientations. We first test whether they can signal deviance for the perpendicular grating stimulus *y* (Figure 8A, B). Both pPE*_x_*and nPE*_y_* respond to stimulus *y* consistently across contexts, supporting the theory that pPE*_x_*selectively responds to S*_x_ −* P*_x_* but not S*_y_*, and that nPE*_y_* neurons receive excessive inhibition from stimulus *y*, consistent with their response to P*_y_ −* S*_y_*.

Furthermore, nPE*_y_* neurons cannot be interpreted as predictor neurons under cPC, whose responses accumulate over repeated presentations of the corresponding stimulus. In cPC, a putative predictor neuron for stimulus *y* should respond similarly to D*_x_* and R*_y_*, while a predictor for stimulus *x* should show a stronger response to R*_x_* than to C*_x_*. Neither pattern matches the observed responses of nPE*_y_* neurons (Figure 7E, Figure 8B).

We then examine orientation selectivity using responses from the many-standard control condition (Figure 8C, D). We find that pPE*_x_*neurons exhibit greater orientation selectivity (Figure 8C), consistent with findings from the motion-visual mismatch experiment [59]. In contrast, nPE*_y_* neurons show weaker selectivity across orientations (Figure 8D). This difference is further supported by measures of orientation selectivity, orientation variability, and average power (Figure 8E), all of which are significantly higher for pPE*_x_*than for nPE*_y_* neurons.

Taken together, these results show that pPE neurons are selective to stimulus orientation, whereas nPE neurons are not.

## Discussion

Here, we demonstrate that both positive (pPE) and negative (nPE) prediction errors, unified under the duet predictive coding (dPC) framework, are essential to predictive coding and are supported by experimental evidence. Within this framework, diverse forms of deviance detection, including omission responses, repetition suppression, mismatch negativity, and oddball responses, arise through the cooperation between pPE and nPE populations. Guided by this theory, we identified putative nPE neurons in oddball paradigm data that had not been previously acknowledged. Moreover, we show that pPE and nPE neurons differ in their orientation selectivity, a distinction that may serve as a functional signature in future experiments. Finally, we find that the strength of deviant responses decrease with increasing similarity between standard and deviant stimuli, as well as with longer interstimulus intervals, a modeling prediction that can be directly tested.

The pioneering work of Rao and Ballard [38] shifted the prevailing view of the sensory cortices from a passive information-processing machine to an active organ of inference [11, 22]. In their formulation, top-down input represents predictions about the external world. Incoming sensory signals are compared against these internal predictions, and only the residual difference—i.e., the prediction error—is propagated to higher levels of the cortical hierarchy. However, the classical predictive coding (cPC) framework and its extensions [32, 47, 31, 12] focus on the pPE component, although the possibility of separate populations for signaling pPE and nPE is occasionally mentioned [38, 32]. Because pPE neurons are suppressed by top-down input, while corticocortical feedback can be net excitatory [58, 10, 23, 57], this mismatch has led to confusion and is frequently referred to as a challenge to the predictive coding framework [11, 19, 12, 32].

Our work suggests that duet predictive coding (dPC) can address limitations of previous models by incorporating negative prediction error (nPE) neurons into the circuit. Inspired by recent experimental findings [1, 25] and computational models [20], we previously argued that a biologically plausible local learning rule can give rise to optimal connectivity for computing prediction errors [30]. This learned connectivity can further explain experimental observations such as motion-modulated suppression and motor–visual mismatch in the context of corollary discharge [30]. In our model, neurons exhibit low baseline firing rates, from which they can be robustly excited but only weakly inhibited. Consequently, top-down input is inhibitory only when paired with an expected stimulus but becomes excitatory in other contexts (Figure 1E). In other words, top-down inhibition is context-dependent rather than absolute.

Under this dPC framework, the deviant response in oddball protocols can be conceptually divided into two components: reduced top-down inhibition to the unexpected stimulus (stimulus *x* in the sequence *yyyx*) and top-down excitation to the omitted stimulus (no stimulus *y* during deviant in the sequence *yyyx*). These effects are mediated respectively by pPE neurons tuned to the unexpected stimulus and nPE neurons selective to the predictor associated with the omitted stimulus. Based on this theory, we identified putative nPE neurons that contribute to deviance detection in the oddball paradigm (Figure 7) and further predict that these nPE neurons will become more clearly distinguishable when combined with an omission protocol (Figure 5).

In our model, the predictors are represented as mutually inhibited leaky integrators with long time constants. A time constant longer than the interstimulus interval is essential for building and maintaining the internal expectation across multiple stimulus repetitions (Figure 5F). Indeed, the time constants in the higher hierarchy areas, such as the prefrontal cortex, are much longer than those in the sensory areas [6, 33, 54]. Other slow mechanisms, including short-term plasticity (STP), may also help maintain internal expectations. However, it remains unclear how STP-related processes could implement the reduction of top-down inhibition compared to the baseline. In our model, the baseline firing rates of predictors are around 5 Hz, higher than those typically observed in sensory areas. Supporting that, the prefrontal cortex is observed to have a higher baseline firing rate [45, 21] than sensory areas, especially in the superficial layer 2/3 [8, 34, 39]. This suggests that a single predictor neuron at a higher hierarchical region can signal the expectancy change by increasing or decreasing its firing rate. However, this is not the only possibility. Theoretically, different pools of neurons can indicate an increase or decrease in internal expectancy as we split the prediction error neurons into pPE and nPE.

Furthermore, the predictors may have attractor dynamics instead of leaky integrators. Currently, we choose the leaky integrator because the error signal decays away with longer interstimulus intervals in a passive task [27]. However, the predictions may be generated from attractor dynamics, especially when the internal prediction must be held to perform an active task, like delayed-match-to-sample tasks [42, 56]. In addition, the internal prediction may be coded through synaptic mechanisms [49, 36]. Finally, as we argued in our previous work [30], internal predictions may be generated *ad hoc*, as in the cases of corollary discharge. For example, when riding a bicycle on a rocky mountain trail, internal predictions of the visual field are continuously updated to account for self-motion and environmental changes. In such situations, precise predictive representations may be transient and do not require maintenance over seconds. In summary, the detailed mechanisms of prediction generation require further study, and they are likely to differ across contexts.

If predictions are generated through different mechanisms in distinct brain regions, are pPE and nPE neurons drawn from the same neuronal pool across experiments? Our previous work [30] argued that the learned pPE or nPE identity of each neuron is determined by the initial ratio of excitatory input from stimulus versus prediction. If the prediction input arises solely from a corollary discharge signal, then pPE and nPE identities should remain stable as long as predictions map consistently onto the same motion signal. In the current study, the stimulus source remains constant, but the prediction source may vary. Since pPE neurons require strong stimulus input, we argue that the pPE population should be conserved across protocols. This is also consistent with the fact that the pPE neurons were strongly orientation selective (Supplementary Figure 8). However, if the prediction source differs, the corresponding nPE population may shift. In our model, we treat variability in input strength as heterogeneity, but recent work suggests that transcriptomic identity may also shape error signaling: specifically, L2/3 *Rrad* or *Baz1a* pyramidal neurons are enriched for pPE responses, while *Adamts2* neurons preferentially exhibit nPE responses [35, 7]. These findings raise the possibility that pPE and nPE neurons may be further distinguished by their molecular and transcriptomic profiles.

Similarly, what are the identities of the two interneuron populations in our model? From a modeling perspective, the simplest approach is to assume an excitation–inhibition balance at both the soma and dendrites [20] or to posit a division of labor across interneuron subtypes [53]. Based on these assumptions, dendrite-targeting somatostatin-positive (SST) interneurons (INs) are likely to mediate stimulus-driven inhibition, while soma-targeting parvalbumin-positive (PV) interneurons are well-positioned to relay prediction-related inhibition [51]. The role of SST cells in delivering stimulus-related inhibition to nPE neurons is supported by studies on motion–visual mismatch [1, 55, 25]. However, the identity of IN subtypes responsible for relaying top-down prediction remains less clear. In the auditory cortex, motion-related inhibition has been attributed to PV interneurons [40, 41]. Yet, recent findings suggest that this pathway may reflect a homogeneous gain control mechanism rather than balancing excitation from a specific predicted stimulus [3]. Although SST cells are generally less directly activated by long-range cortical inputs [29], emerging evidence indicates that they may, in some cases, be more responsive than PV cells in certain top-down signaling pathways [43]. Their involvement in pPE calculation is also supported in both auditory [41, 3] and visual studies [16]. Another major interneuron subtype, vasoactive intestinal peptide-positive (VIP) cells, typically inhibit SST cells and disinhibit pyramidal neurons [37, 58], making them less likely to mediate top-down inhibition directly. However, our recent findings suggest that a subpopulation of VIP interneurons co-expressing the neuropeptide cholecystokinin (CCK) can directly inhibit pyramidal cells with synaptic strength comparable to that onto SST cells, implying a potential role in mediating top-down inhibition [9]. In summary, the identity of interneurons responsible for relaying top-down inhibition remains unresolved. Given that internal predictions can arise via diverse mechanisms and pathways, it is plausible that top-down inhibition is mediated by a combination of IN subtypes rather than a single, dedicated class.

Classical predictive coding (cPC), with its focus on top-down inhibition, fails to account for many experimental observations. Here, we propose duet predictive coding (dPC), which incorporates both positive-prediction error (pPE) and negative-prediction error (nPE) neurons to bridge the gap between theory and diverse experimental findings. Our framework reveals context-dependent top-down modulation of sensory processing across multiple predictive coding paradigms, offering new insight into the generality of predictive coding. We further identify behavioral signatures that differentiate nPE from pPE neurons, opening new avenues for experimental investigation.

## Methods

### Sensory area

In this work, the single-neuron and synapse models within the sensory area are the same as in our previous study [30]. The connectivity within each column also follows the learned connectivity from [30]. The neuron index within each column is sorted based on the input strength of the preferred stimulus. When showing the example traces, the pPE example neuron is always the neuron with the smallest index (maximum stimulus input strength), and the nPE example neuron is the one with the largest index (minimum stimulus input strength).

We employ two versions of the model in this study. The simplified version assumes homeostasis, meaning that the total maximum input from stimulus and prediction is equal across neurons. This version includes a small number of neurons per column (*N_pCol_* = 40) and is used throughout most of the paper. It offers a more interpretable framework for illustrating the core mechanisms of dPC. The realistic version, in contrast, is designed to match experimental statistics and render testable predictions from theory. This version includes more neurons per column (*N_pCol_* = 200), and it does not assume homeostasis. While it is less intuitive for illustrating mechanisms, it is essential for capturing biological variability, especially since homeostasis is not guaranteed *in vivo*, as seen in studies of selective modulation of auditory responses [3, 2]. Throughout the paper, we use the simplified version by default, and results from the realistic version are always explicitly stated.

In the following sections, we adopt the same notations as in [30]. In addition, we introduce predictors that generate top-down prediction inputs, and we extend the model to include multiple columns with different orientation selectivity. The specific modifications are described below:

### Predictor

The pyramidal cells within a column are connected to an integrator with the same preferred orientation and homogeneous connectivity strength *g_Int,E_*. These integrators are modeled as

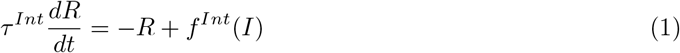

where *τ ^Int^* has a much longer time constant. The activation function here is a saturated version of the previous activation function *f* :

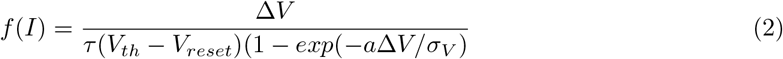

with Δ*V* = *I/g_L_* + *V_l_ − V_th_* and the saturation as follows:

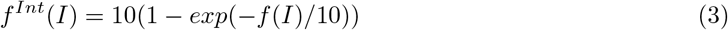

With this choice, the activity of the integrator is capped at 10Hz, which is the maximum firing rate for the predictor in the previous model [30].

The integrators mutually inhibit each other through a bell-shaped connectivity.

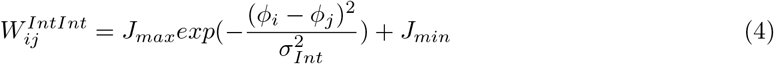

Here, *J_min_* is negative to represent lateral inhibition. For simplicity, we don’t split recurrent excitation and inhibition in different connectivity matrices.

### Simulated stimulus input

The stimulus input to individual neurons within the same column is drawn from a linearly uniform random distribution ranging from zero to a maximum value 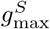, as in our previous work [30]. In simulations involving multiple columns, the maximum input is scaled based on the angular difference between a neuron’s preferred orientation (frequency) 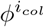 and the stimulus orientation (frequency) *ϕ*^0^, using a Gaussian kernel:

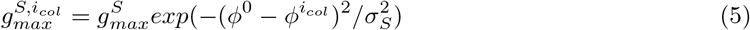

With *σ_s_* is the width of the Gaussian kernel.

In the fixed oddball protocol, we simulate the model with stimulus *S_x_* (standard stimulus), presented at a fixed orientation for eight repetitions. Each stimulus is shown for a duration *t_S_*, with an inter-stimulus interval of *t_Int_*. Following the eighth repetition, we present an orthogonal stimulus *S_y_* as the deviant. This sequence is repeated twice and then followed by two additional standard trials with *S_x_*. In the omission protocol, the deviant stimulus is replaced with an omission (i.e., *S_y_* = 0). In the randomized oddball protocol, the stimulus on each trial is determined probabilistically based on the assigned probabilities *P_x_* and *P_y_*.

In the fixed oddball and omission protocols, adaptation is modeled by directly reducing themaximum stimulus conductance 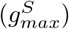 for repeated stimuli.

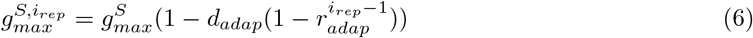

In the simulation matching MMN (Figure 3), adaptation is set to zero (*d*_adap_ = 0), based on lesion study findings [26], and is also omitted in the randomized oddball sequences.

### Top-down prediction input

In our model, we include a prefrontal cortex (PFC)-like circuit composed of leaky integrators that accumulate temporal information. We assume that each predictor *j* projects only to its corresponding sensory column, generating mismatch receptive fields that match that of the sensory receptive field [59]. The top-down connectivity is the same as the learned connectivity matrix *W_ij_* in our previous work [30], where *j* indexes the *j*-th predictor and *i* belongs to the *j*-th column.

These predictors excite the dendrites of pyramidal cells as well as the interneuron population *I*_2_. Experimental evidence suggests that top-down predictions may exert their influence with temporal precision, such that the sensory area receives prediction input only when it is computationally relevant for local processing. This timed prediction input could be implemented biologically via spike-timingdependent plasticity (STDP), as proposed in [52]. For simplicity, we implement it as a binary timing in our model: the top-down excitation deviating from baseline is only applied when a stimulus is either presented or expected.

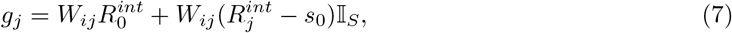

𝕀*_S_* = 1 when any stimulus is presented. In the omission protocol, we also set I*_S_* = 1 when the model is “expecting” a stimulus.

### Data analysis

In the auditory omission response analysis (Figure 2), we use the same omission index to quantify the distribution of population responses as in [27]. Specifically, for each cell, let *r_s_* denote the response to the last stimulus presentation, and *r_o_* the response to the omission. The omission index is defined as: iOMI = (*r_o_*−*r_s_*)*/*(*r_o_* + *r_s_*). In our model, omission-responsive neurons are defined as those whose response during the omission period is at least twice as large as during the pre-omission period.

In reproducing MMN (Figure 3), we used a similar method to calculate “memory-based expectancy” *M_s_* as in [48]: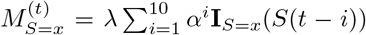, with **I***_s_* is an identity function where **I**_*S*=*x*_(*x*) = 1 and **I**_*S*=*x*_(*y*) = 0, and the normalization factor 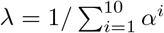.

In analyzing pPE and nPE population responses in the model, a neuron is labeled as a pPE neuron if it shows a positive deviance detection and a negative omission difference. Conversely, it is labeled as an nPE neuron if both the deviance detection and omission difference are positive. After classification, activity is averaged separately across the pPE and nPE populations.

To test whether nPE neurons play a functional role in the oddball paradigm, we analyzed experimental data from mice of either sex (*n* = 3), including previously published recordings from [5].

Detailed procedures for data collection and z-score normalization are described in [5]. The total number of cell–stimulus pairs examined was 326. A cell was classified as responsive only if its response to a stimulus in the many-standard control condition was significantly greater than its pre-stimulus baseline (*p <* 0.05). We then focused on the subset of responsive cells that showed significantly higher activity in response to the same stimulus in the deviant context compared to the many-standard control (*p <* 0.05). All *p*-values for single-cell comparisons were corrected using the Benjamini–Hochberg (BH) procedure.

## Code Availability

We analyze the model-generated data using Python 3.9 and the experimental data using MATLAB R2024b. All analysis code will be made available on GitHub upon acceptance.

## Declaration of generative AI and AI-assisted technologies in the writing process

During the preparation of this work, the authors used ChatGPT-4o in order to assist with proofreading and improving the clarity of the manuscript. After using this tool, the authors reviewed and edited the content as needed and take full responsibility for the content of the publication.

## Author contributions

J.H.M. and X.J.W. conceptualized the project. J.H.M. performed simulations, data analyses, and wrote the manuscript. J.M.R. and J.P.H. collect the sample. J.P.H. and X.J.W. supervised the study. All authors contributed to editing and finalizing the manuscript.

## Acknowledgement

We thank Aldo Battista, Yue Liu, Tianshu Li, and other members of Xiao-Jing Wang lab for the discussion. We thank Cristina Savin, David Schneider and Georg Keller for their suggestions and feedback on an early version of the manuscript. We thank Kevin Berlemont for an early exploration of this project. This work is supported by ONR grant N00014-23-1-2040 and NIH grant R01MH062349 (to XJW), and NIH grant R01EY033950 from the National Eye Institute (to JPH)

## Supplementary Materials

